# Flexible Fitting of Biomolecular Structures to Atomic Force Microscopy Images via Biased Molecular Simulations

**DOI:** 10.1101/793901

**Authors:** Toru Niina, Sotaro Fuchigami, Shoji Takada

**Affiliations:** Department of Biophysics, Graduate School of Science, Kyoto University, Kyoto 606-8502, Japan

**Keywords:** coarse-grained molecular dynamics, HS-AFM, afmize, CafeMol

## Abstract

Atomic force microscopy (AFM) is a prominent imaging technology that observes large-scale structural dynamics of biomolecules near the physiological condition, but the AFM data are limited to the surface shape of specimens. Rigid-body fitting methods were developed to obtain molecular structures that fit to an AFM image, without accounting for conformational changes. Here we developed a method to fit flexibly a three-dimensional biomolecular structure into an AFM image. First, we describe a method to produce a pseudo-AFM image from a given three-dimensional structure in a differentiable form. Then, using a correlation function between the experimental AFM image and the computational pseudo-AFM image, we developed a flexible fitting molecular dynamics (MD) simulation method, by which we obtain protein structures that well fit to the given AFM image. We first test it with a twin-experiment; for a synthetic AFM image produced from a protein structure different from its native conformation, the flexible fitting MD simulations sampled those that fit well the AFM image. Then, parameter dependence in the protocol is discussed. Finally, we applied the method to a real experimental AFM image for a flagellar protein FlhA, demonstrating its applicability. We also test the rigid-body fitting of a fixed structure to the AFM image. Our method will be a general tool for structure modeling based on AFM images and is publicly available through CafeMol software.

## INTRODUCTION

Elaborate mechanisms of cells are realized by functional biomolecules that often undergo conformational changes to show their functionalities. Resolving their conformations at work is an essential step to understand the underlying physical machinery of life. However, it is still difficult to observe the process of conformational changes of working biomolecules directly in near atomic resolution. While X-ray crystallography is a powerful method that has the ability to determine the atomic coordinates^1^, it requires crystallization of the protein stabilizing specific conformational states. Recent revolution in cryo-electron microscopy (cryo-EM) enables to obtain structures at atomic resolution without crystallization^2,3^. However, in order to gain high resolution, cryo-EM requires freezing into ultra-low temperature and averaging images for individual conformations. Thus, while at high-spatial resolution, X-ray crystallography and cryo-EM provide snapshots of biomolecules at somewhat non-physiological condition.

The atomic force microscopy (AFM) is a prominent microscopic method that is used extensively to acquire high-resolution images directly under near physiological conditions in the field of molecular biology^4–7^. Unlike other microscopes, it scans the specimen with a physical probe that monitors the force from the surface of molecules. Owing to the efforts to increase the frame acquisition rate, nowadays a high-speed AFM (HS-AFM) takes tens of frames per second^7^. Typically, HS-AFM achieves ~2 nm lateral resolution and ~0.15 nm vertical resolution^7^. Importantly, the measurement process is gentle enough to preserve the structures and functions of biomolecules. It is said that every tapping by the vibrating cantilever tip gives 2 to 5 kBT on average to its specimen^5^. Additionally, the energy provided by tapping dissipates immediately into several degrees of freedom of the system including water molecules. Thanks to its spatiotemporal resolution, HS-AFM is employed to capture the dynamics of working biomolecules, such as F_1_ ATPase^8^, Myosin V^9^, bacteriorhodopsin^10^, CRISPR-Cas9^11^, and many others and provided us new insights about their structures at work. Low-invasive observation of HS-AFM enables to capture the dynamics of working biomolecules at the single molecule resolution.

Although the HS-AFM is widely used to investigate the dynamics of functional biomolecules, there is no tool to reconstruct the structure from AFM image through flexible fitting method. The resolution of AFM images is not high enough to construct an atomic structure model from scratch. Therefore, high-resolution structures of the molecules obtained by other experimental observations are useful as the starting point to reconstruct three-dimensional molecular models. Generally, the analysis begins with detection of image areas in which the molecules of interest are located; there are several methods that are utilized for that purpose, such as binary digitization^12^ and volumetric analysis^13^. As a next step, for large molecules and macromolecular complexes, there are methods to perform rigid-body fitting to determine the position and orientation of each protein or its domains that is optimal by certain criteria^14–16^. Those methods were applied not only to macromolecular complex structures^17,18^, but also to the arrangement of domains of a large protein^19,20^ and successfully determined the relative orientation. Though the methods have an ability to determine the most likely arrangements of proteins and domains, the rigid body fitting methods have a limitation when investigating conformational changes of molecules because these methods treat structures of the proteins/domains fixed in their known structures.

In this work, we developed a new method to obtain structural information from AFM images by integrating a cost function that represents similarity between structural models and AFM images with molecular dynamics (MD) simulations. To quantitatively compare the structural models and AFM images, we first convert a three-dimensional structural model into a pseudo-AFM image. Since the previous algorithm to generate pseudo-AFM image from structural models is not applicable to MD simulations due to non-differentiability, we first developed an algorithm to generate a pseudo-AFM image from a structure model in a differentiable form, which serves as a cost function. Next, we applied the cost function to the rigid-body Monte-Carlo simulation and confirmed that our method successfully found the appropriate orientation from noisy AFM images by minimizing the cost. After that, using our new cost function, we constructed a new force field that enables flexible fitting into AFM images. We combined our biasing potential, i.e., the cost function, with a well-established coarse-grained protein model, AICG2+^21–23^, and applied it to several test proteins with varying structural characteristics. In a twin-experiment, by comparing the resulting conformations and the reference structure that is used to generate target AFM image, we confirmed that our new flexible fitting method successfully samples near-optimal structures. Lastly, we demonstrated that our method works with real experimental data. Starting from the crystal structure placed by hand, our method successfully increases the similarity between the experimental AFM image and the structural model, giving plausible three-dimensional structures that are compatible with the experiment.

## Methods

### Collision-detection pseudo-AFM Image Generation

We begin with a conventionally used method to generate a pseudo-AFM image for a given protein structure model (Figure 1A). This method will be used to quantify the fitness of a structural model with the experimental AFM image; we will use a correlation between the experimental AFM image with the pseudo-AFM image generated from the structure model. Previously, a method has been used to compare experimentally obtained AFM images with pseudo-AFM images generated from known PDB structures^8^. This conventional method generates the height information corresponding to each pixel by calculating a steric collision between the tip of a cantilever and atoms in the molecular models by moving tip position down (Figure 1B). As the tip of cantilever, we assume a shape composed of a cone capped by a sphere and choose r=1.0 nm for the tip radius and θ = 5 degree for the half apex angle. Then, we can calculate the height for a given molecular structure. In this work, we took this approach, termed the collision-detection method, to generate a reference pseudo-AFM image. The collision-detection method is available as a tool “afmize” at http://doi.org/10.5281/zenode.3362044.

**Figure 1.**
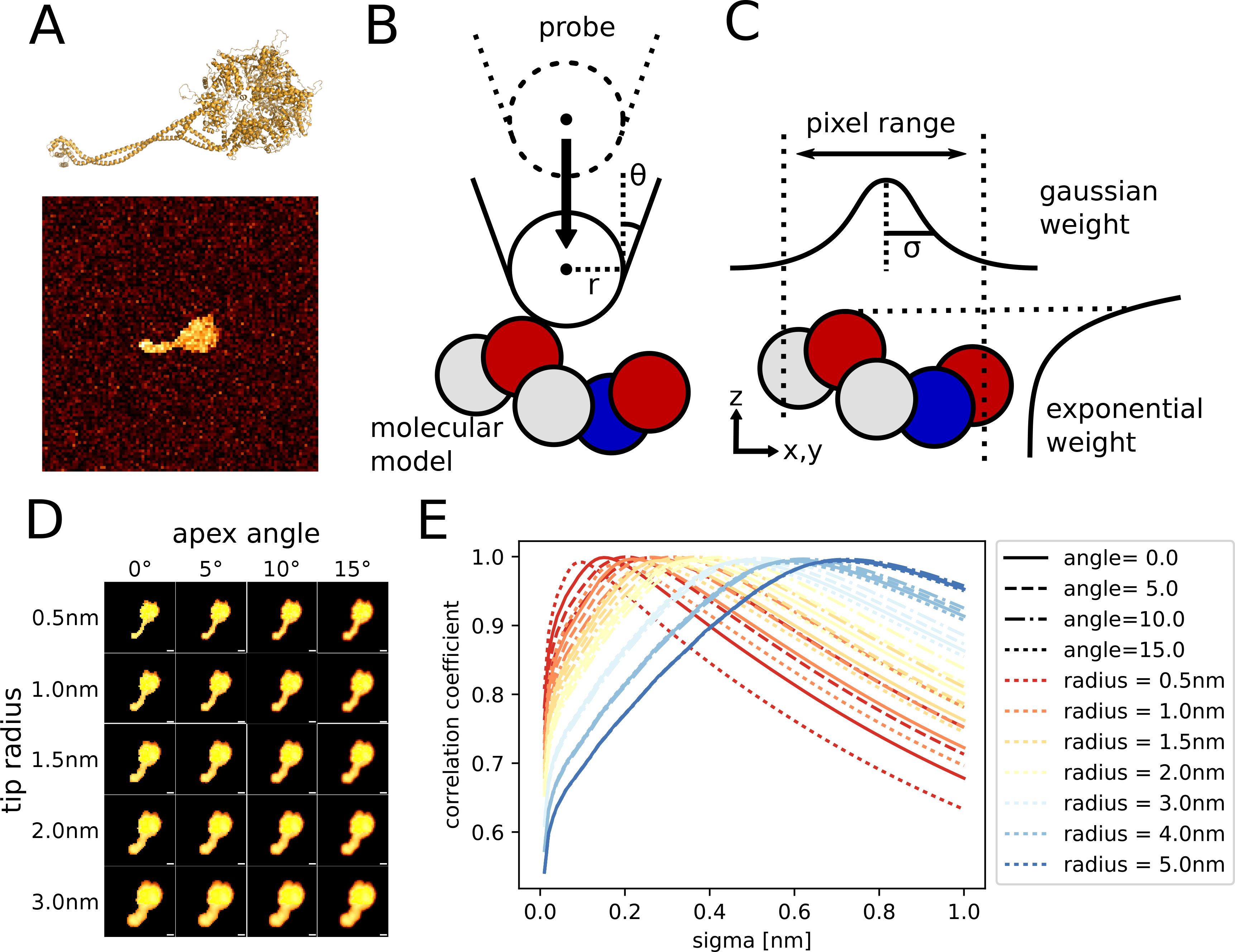
(A) The structure of a test protein, dynein (upper) and an example of its synthetic AFM image constructed by the collision-detection method with a noise (lower). (B) A cartoon for the collision-detection method in “afmize” to generate the pseudo-AFM image by detecting the collision between the cantilever tip and a molecular model. *r* and *θ* represent the tip radius and the half of apex angle, respectively. (C) A cartoon for the method to generate a smoothed AFM image. (D) Pseudo-AFM images of dynein generated by the collision-detection method with varying parameters *r* and *θ*. As the parameters increase, the images become more blurred. (E) The correlation between the collision-detection and the smoothed pseudo-AFM images. The radius and angle used are indicated by the color and line type of every curves, respectively. By changing the value of *σ*, the smoothed image-generation method can achieve sufficiently high correlation with the one generated by the collision-detection method.

### Smoothed Pseudo-AFM Image Function

Since the above procedure cannot be described as a smooth and differentiable function, the procedure is not directly applicable as a potential energy function in flexible-fitting MD simulation. Hence, we propose a new method to generate pseudo-AFM images in a differentiable form, approximating the above method by a set of smooth weighting functions.

For a target biomolecule represented by its coordinate (*x*_*j*_, *y*_*j*_, *z*_*j*_) and radius *r*_*j*_ for the *j*-th particle (*j* = 1, …, *N*), we define a height function 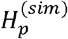 at the *p*-th pixel (*x*_*p*_,*y*_*p*_) as

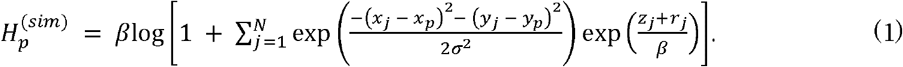

Here, without loss of generality, we define xy-plane as the stage of the AFM so that the z-coordinate represents the height of the specimen. In this function, we used two weighting functions to each particle. One is a Gaussian function which measures the closeness between the *p*-th pixel center and the *j*-th particle in *xy*plane and takes a value close to 1 when a particle locates near the center position of the *p*-th pixel. The parameter *σ* represents the spatial range. The other weighting function is the so-called logarithm-summation-exponential function of the position along the z coordinate of each particle, which is a differentiable approximation to the maximum function. *β* is a parameter that makes the function smooth; the larger *β* is, the smoother the surface 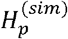 is. The constant 1 in the logarithm function represents the stage height at the absence of the specimen.

In this work, the *β* is set to 0.1 nm, which is the smallest value where the numerical calculation is still stable with our target proteins in this paper. With even a smaller *β*, the numerical errors become intolerable and make the whole process unstable. The *σ* s are determined by comparing the image generated by the proposed method and the traditional method. We chose the value that makes the highest correlation between the images generated by traditional method and our smoothed method (see “Results”).

### Stage Potential

To observe biomolecules by AFM, target molecules are placed on the AFM stage ^[citation-needed]^. We set the planar AFM stage as the plane z = 0. We modeled the interaction between the AFM stage and biomolecules as a simple Lennard-Jones potential along z axis,

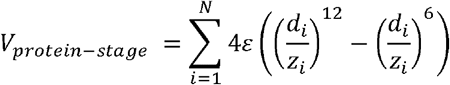

where *d*_*i*_ is the radius of *i*-th particle. We choose *ε* = 1.2 kcal/mol that is weak enough to allow them to diffuse and rotate on the surface without breaking its structure.

### Biasing Potential

To achieve the maximum similarity between a pseudo-AFM image generated from a structure model and the experimental AFM data, we introduce a biasing potential (a cost function) that is a differentiable function of coordinates and thus that can be used in MD simulations. For the functional form, we employed a modified correlation coefficient *c*.*c*.(***R***) to define the form of the potential, motivated by^18^, defined as

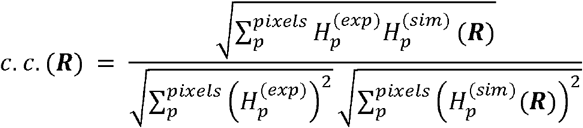

The biasing potential is defined by

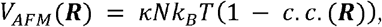

where *k*_*B*_ is the Boltzmann constant, *T* is the temperature, *κ* is a dimensionless parameter that controls the strength of the bias. The appropriate value of *κ* will be discussed later.

### Protein Modeling

We used two proteins, a fragment of tight-junction protein ZO-1 (PDB ID: 3SHW) and the cytoplasmic domain of a flagellar protein FlhA (3A5I) ^26,27^ as target molecules in our test cases. Missing residues and atoms in the middle of PDB structures were modeled by Modeller^28^. We simulated all the target proteins using the AICG2+ model where each amino acid is represented by a coarse-grained bead located at the position of corresponding Cα atom. Because ZO-1 and PDZ12 have some flexible linker region, we removed native contacts between linker and globular domains and native contacts between globular domains.

We also used dynein ^29^ and hemoglobin ^30^ in our rigid-body Monte-Carlo simulations. For them, we used all the heavy atoms in the PDB structure. Since these are modeled as a rigid body, we just keep their structure during the whole simulation processes.

### Rigid Body Metropolis Monte-Carlo Simulation

For comparison, we performed Metropolis Monte-Carlo simulated annealing for the rigid-body fitting to the AFM image. Method details are presented in the Supporting Information.

### Flexible Fitting Molecular Dynamics Simulations

In the flexible fitting MD simulations, the total energy function, *V*_*total*_, is given as a sum of the protein force-field, *V*_*protein*_, the stage potential, *V*_*protein-stage*_, and the biasing potentials towards the AFM image, *V*_*AFM*_;

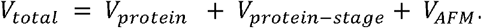

The simulations were conducted by Langevin dynamics at a constant temperature of 300K for 10^8^ MD steps. All the simulations are conducted by an in-house program named Mjolnir that is freely available on GitHub; https://github.com/Mjolnir-MD/Mjolnir. The tool is implemented in CafeMol^31^ as well, which is also available through its website (www.cafemol.org).

## RESULTS

### Verification of Smoothed AFM-image generation method

We first verified that a smoothed pseudo-AFM image generated by out new approach is close enough to a pseudo-AFM image obtained by the collision-detection method. As the target protein, we chose the structure of dynein (Figure 1A) because it is large and has several structural elements and thus can be a good test structure. First, we generated pseudo-AFM images using the collision-detection method with several tip sizes and apex angles as reference images (Figure 1D). Then, we generated the smoothed pseudo-AFM images with varying σ values. The similarity between these two AFM images were quantified as *c*.*c*.(***R***), the same metric we use for the flexible fitting MD later. Figure 1E shows correlations between results obtained by two methods. For any combination of two parameters *r* and *θ* tested, we can achieve the correlation value over 0.98 by choosing the appropriate *σ* value (Table 1). It means that by using our smoothed pseudo-AFM image generation algorithm, we can generate almost the same pseudo-AFM image having high correlation with an image obtained by conventional collision-detection method.

**Table 1.**

|  |  |  |  |  |  |  |  |
| --- | --- | --- | --- | --- | --- | --- | --- |
|  | 0.5 [nm] | 1.0 [nm] | 1.5 [nm] | 2.0 [nm] | 3.0 [nm] | 4.0 [nm] | 5.0 [nm] |
| 0.0 [deg] | 0.09 | 0.18 | 0.26 | 0.32 | 0.47 | 0.59 | 0.71 |
| 5.0 [deg] | 0.15 | 0.22 | 0.28 | 0.35 | 0.48 | 0.6 | 0.71 |
| 10.0 [deg] | 0.2 | 0.26 | 0.32 | 0.38 | 0.5 | 0.61 | 0.72 |
| 15.0 [deg] | 0.27 | 0.31 | 0.37 | 0.42 | 0.52 | 0.62 | 0.72 |

### Rigid Body Metropolis Monte-Carlo Simulation

Before starting flexible fitting MD simulation, using – *c*.*c*.(***R***) as the cost function, we performed the rigid body Metropolis Monte-Carlo simulations with the simulated annealing as a simpler case to confirm whether our cost function works successfully.

First, we used dynein consisted of multiple structural units as our test protein because it is large enough so that most of the structural units can be distinguished in the AFM image and it has an asymmetric structure that can be considered to be easy to find the correct orientation. We first generated a pseudo-AFM image by the collision-detection method of randomly rotated protein structure and added Gaussian noise to the image. Then, we used this pseudo-AFM image as the reference (synthetic) AFM image in order to evaluate the correctness of results. Starting from a randomly-oriented initial structure, we performed rigid body Metropolis Monte-Carlo simulation (Figure 2A) to maximize *c*.*c*.(***R***). To compare the reference structure and the snapshot during the simulation, we used the root mean square displacement (RMSD) without aligning them (Since we used the rigid body model, if we align it, the RMSD value becomes zero). In most of the Monte Carlo runs, as the number of steps increases with the temperature decreasing, correlation coefficient increases and the RMSD value decreases (Figure 2B). Thus, our rigid-body approach can find the location and orientation close to the ground truth image (RMSD = 0.0298 nm) by optimizing the correlation between pseudo-AFM image based on the snapshot and the reference AFM image.

**Figure 2.**
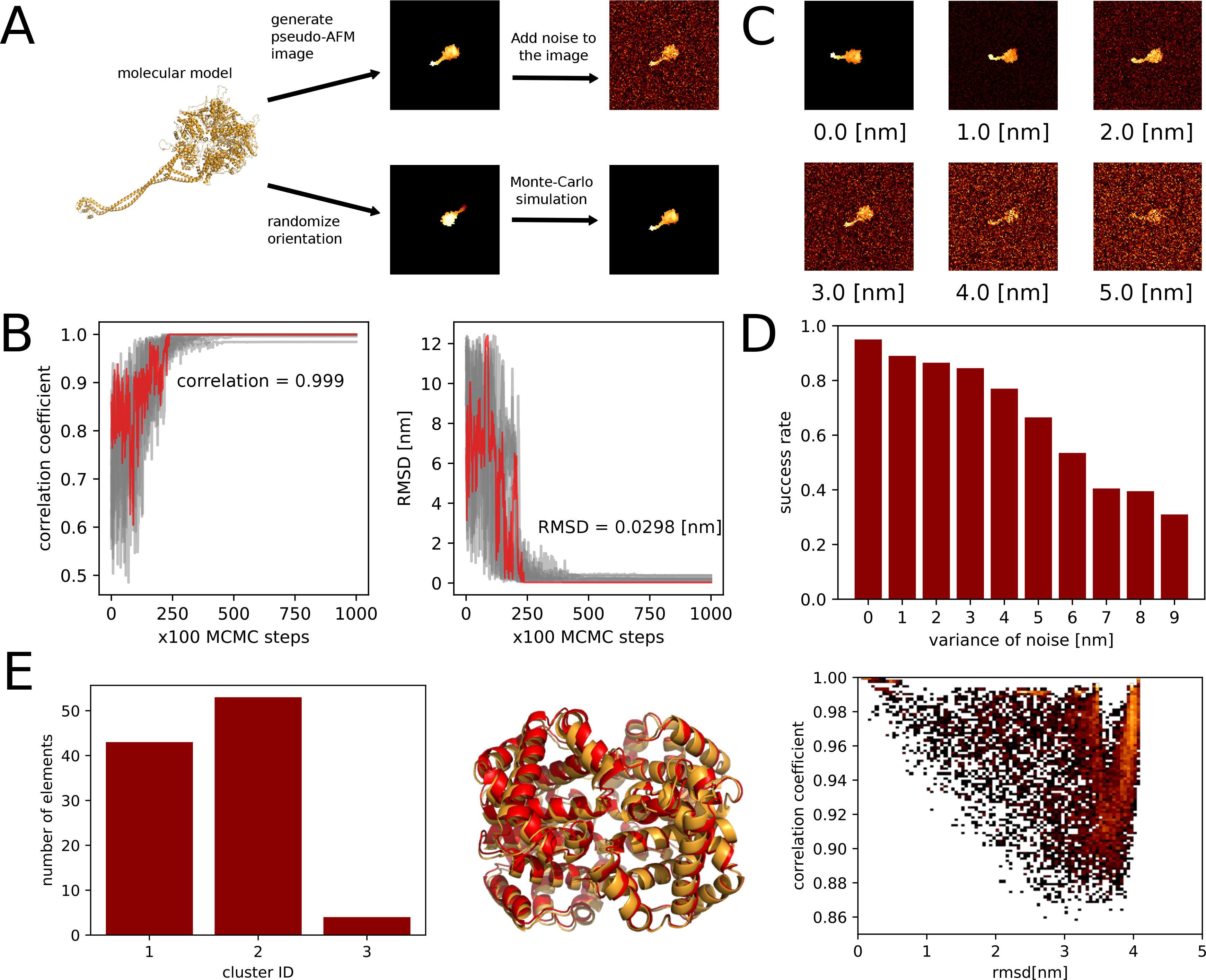
Summary of the rigid-body fitting test of a synthetic AFM image of dynein (A-D) and hemoglobin (E). (A) The outline of the twin experiment protocol. (B) The trajectories of correlation coefficient (left) and RMSD (right) for 10 independent simulated annealing runs. The trajectory that finds the most correlated snapshot is colored red. (C) Pseudo-AFM images on which varying Gaussian noise applied. (D) The number of trajectories that successfully found the correct orientation out of 100 independent simulations with different noise levels. We considered the trajectories found the correct orientation if the final snapshot has RMSD value smaller than 0.5 nm. (E) Histogram of the number of structures in clusters (left), representative structures in two major clusters (red: cluster 1, orange: cluster 2) (middle), and two-dimensional heat map of correlation coefficient and RMSD (right).

Second, to evaluate the robustness of our approach against the noise level, we added spatially-independent Gaussian noise of different levels to our reference pseudo-AFM images (Figure 2C). Then we repeated the same Monte Carlo simulation as above and counted how many simulations find the protein configurations within 0.5 nm of RMSD from the reference structure (Figure 2D). As expected, the success rate decreases as the noise level increases. It becomes difficult to find the correct configuration if the variance of Gaussian noise becomes larger than 3.0 nm in this case.

Next, we performed the same twin experiment for hemoglobin that has a symmetric structure. The hemoglobin is a hetero-tetramer (αβ)_2_ with α and β subunits being homologous and thus possesses the two-fold symmetry and four-fold pseudo-symmetry. We found that most of trajectories fell into one of the two configurations, due to its underlying symmetry. By the k-means clustering on the ensemble of final configurations, we obtained two major and one minor clusters. One corresponds to the same orientation as the reference and the second one corresponds to the symmetry-related and thus equivalent configuration (Figure 2E). The minor third cluster corresponds to the wrong placement caused by local minima in the biasing potential. This third cluster exemplifies potential mis-fitting of the structure when the target molecule contains more than two mutually similar parts.

### Flexible Fitting to the AFM Image using Molecular Dynamics Simulation

Next, we examine whether our biasing potential finds appropriate protein configuration that is far from initial structure, with a twin experiment. To test the capability of our biasing potential in flexible fitting, we used a fragment of tight junction protein, ZO-1 that contains two globular domains that are connected by a flexible linker. First, we performed a short MD simulation to obtain an open configuration of the ZO-1 protein starting from the crystal structure. Next, using the last snapshot of the simulation, we generated a synthetic AFM image using the collision-detection AFM image generation method (Figure 3C the right most image). Using the synthetic AFM image, we performed 40 independent constant temperature MD simulations for flexible fitting.

**Figure 3.**
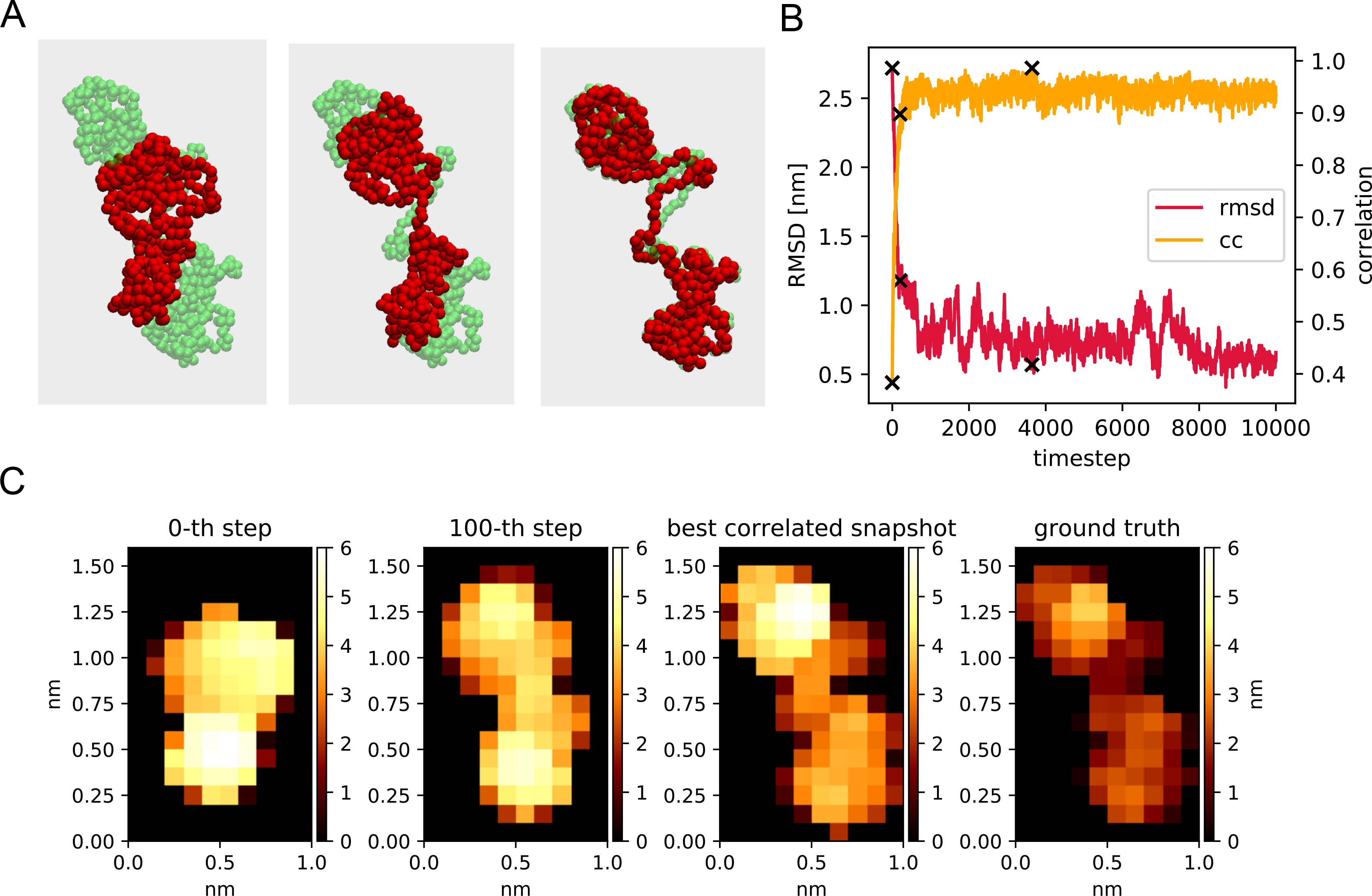
A twin-experiment for the flexible fitting MD simulation to the AFM image for a fragment of tight junction protein, ZO-1. (A) The snapshots in the representative trajectory are drawn in red particles. The ground truth configuration used to generate the synthetic AFM image is shown in green. The first (left) snapshot; the initial configuration. The second; 100-th step snapshot. The last(right) snapshot; the most correlated configuration. (B) The time course of the value of RMSD from the ground truth (red) and the correlation coefficient (yellow). The points that correspond to the shown snapshots are shown in cross markers. (C) The corresponding synthetic AFM images generated from the snapshots shown in A and the reference image.

The representative trajectory shows that our method successfully increases the correlation and decreases RMSD from the ground truth (Figure 3B). Unlike the case of rigid body MC simulations, here we calculated RMSD values after structure alignment. Some of the snapshots during the simulation are shown in Figure 3A and the corresponding AFM images in Figure 3C. It can be seen that the protein approaches toward the reference structure through conformational changes. It means that by optimizing the structure according to our novel biasing potential, the corresponding protein structure can be found.

### Sampling Efficiency and the Appropriate Values of Parameters

The biasing potential has a parameter *κ* that modulates the overall strength of the bias. We expect that a too large value of *κ* leads to unacceptable conformational deformation of proteins and overfitting. Instead, a too small value of *κ* makes the convergence slower and increases the required simulation time. To find the optimal value of *κ*, we performed large-scale conformational sampling with various *κ* values; 40 independent constant temperature MD simulations of 10^8^ MD steps at 300K for each *κ* value.

In the case of *κ* = 0.1, the smallest value tested (Figure 4A), the peak in the population appears at a region with the correlation ~0.6 and the RMSD value ~2.0 nm, which are largely deviated from the ground-truth structure. We, however, note that the sample reaches at the correlation coefficient of 0.915, where the RMSD was 0.763 nm. With this parameter, one needs to sample very long to obtain a reasonably fit structure as a rare sample. Next, Figure 4B shows the result for *κ* = 1, in which the biasing potential strength is comparable to the kinetic energy. The population peak appears around the region where correlation is high (~0.95) and RMSD value is small (~0.7 nm). Importantly, in this case, there is a tendency that the highest correlation corresponds to smaller RMSD values to the ground truth. Using the sample with the highest correlation (0.986), we obtain the structure with its RMSD = 0.569 nm, which is better than the case of *κ* = 0.1. Contrary, in the cases of larger, values (Figure 4C for *κ* = 10, and D for *κ* = 100), there appear many distinct high-correlation structures which includes not only small RMSD snapshots, but also large RMSD structures. Thus, as expected, a strong biasing potential leads structures to deformed states. It seems that the total biasing energy should be comparable to the total kinetic energy, i.e. the value of *κ* should be in the order of 1.

**Figure 4.**
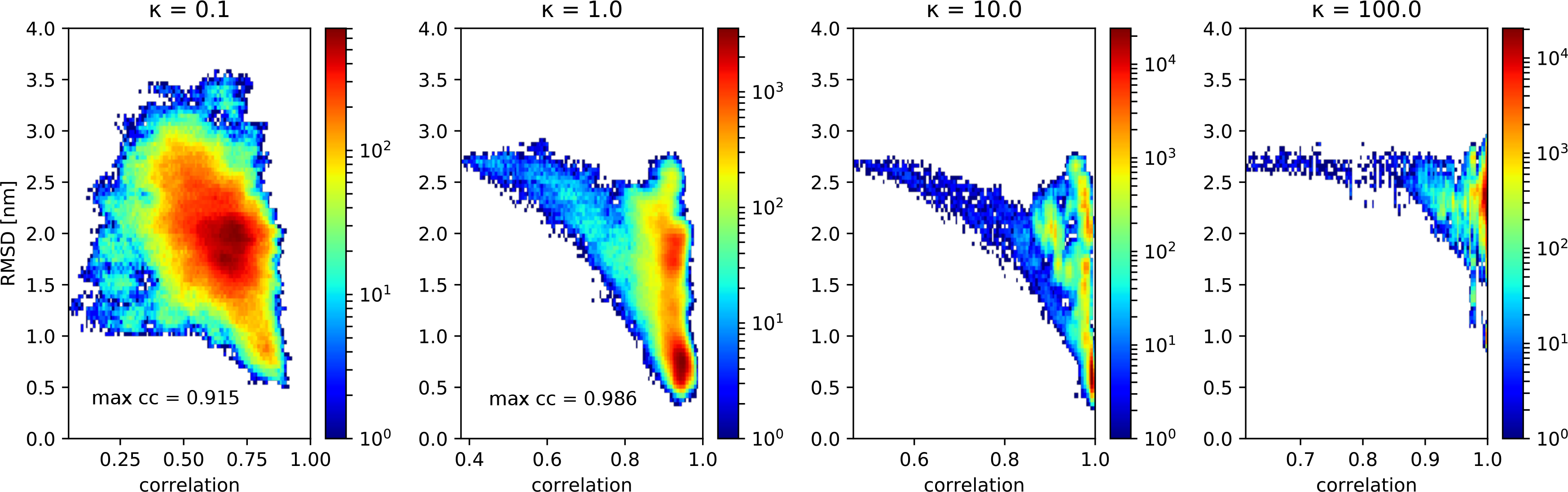
Sampling and test for the κ values for the flexible fitting MD simulation to the AFM image for a fragment of tight junction protein, ZO-1. The κ values, 0.1 (A), 1.0 (B), 10.0 (C), and 100.0 (D) are tested. For the most correlated snapshot, the correlation coefficient and the RMSD from the ground truth were 0.915 and 0.763 nm, respectively, in A and 0.986 and 0.569 nm, respectively, in B.

### Application to an Experimental AFM Image: A Flagellar Protein FlhAc

So far, we used synthetic-AFM images as a reference because the validation of the simulation requires the knowledge of the ground truth; the reference structure. But, it is also necessary to test our method with the real high-speed AFM image.

To test that, we used the experimental data of the cytoplasmic domain of a flagellar protein FlhAC, of which the AFM measurement was recently reported^27^. We choose three snapshots from the high-speed AFM movie of a monomeric FlhAC and took regions in which FlhAc monomer can be identified (Figure 5A, E, I). For each AFM image chosen, we performed 40 independent flexible fitting simulations of 10^8^ MD steps starting from the crystal structure that are placed roughly at the center of the image regions. In all three cases, our simulations found configurations that have higher correlation coefficients (Figure 5B, F, J) than that of the initial configuration.

**Figure 5.**
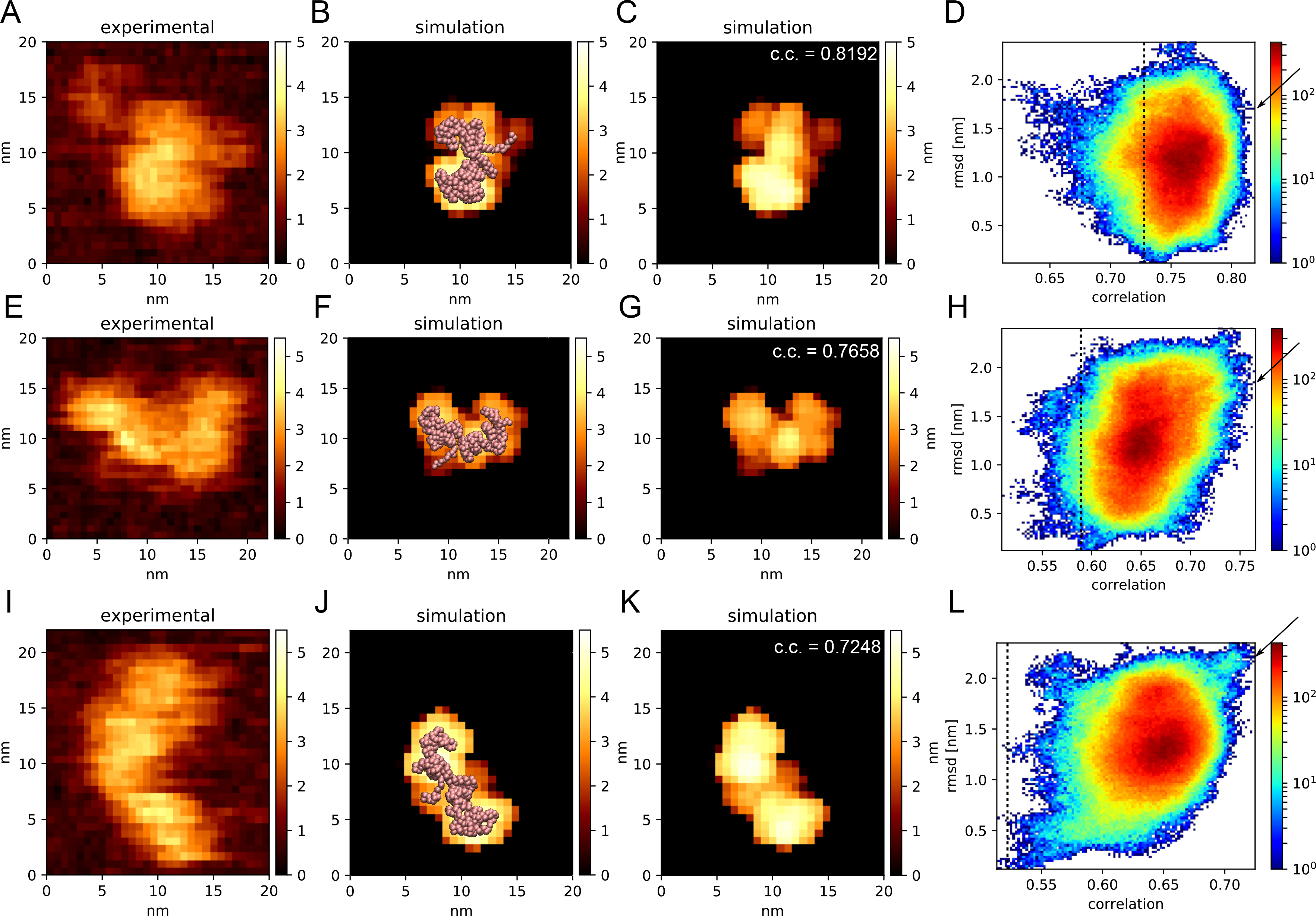
The application of the flexible fitting method to the experimental AFM image. (A, E, I) Portions of experimental snapshots that shows FlhAc molecule. (B, F, J) The snapshots that have the highest correlation value to the experimental image superposed onto the synthetic AFM images. Their RMSD values from the PDB structure were 1.687 nm, 1.836 nm, and 2.197 nm, respectively. The snapshots are drawn using VMD^32^. (C, G, K) Synthetic AFM images generated from the simulation snapshots that have the highest correlation value. (D, H, L) Probability distributions in the simulation. The initial values of correlation are plotted by dotted lines. The arrows point where the most correlated snapshots belong to.

Interestingly, the results of flexible fittings are rather different in the three cases. Of the three images, we obtained a best-fit conformation with the correlation coefficients (CCs), 0.8192 (the largest), 0.7658, and 0.7248 (the smallest), for Figure 5ABC, EFG, and IJK, respectively. This order is anti-correlated with their RMSD values from the crystal structure, 1.687 nm, 1.836 nm, and 2.197 nm for Figure 5ABC, EFG, and IJK. respectively (Note that the RMSD is not from the ground truth, but from the crystal structure, in this case). We also note that for the case of Figure 5A image, the second-best CC = 0.8173 was obtained by a structure with its RMSD = 0.776 nm, which is much closer to the native one. It seems that the larger CC can be achieved when structures relatively close to the native one is compatible. Albeit unclear, we can speculate a plausible reason; structures close to the native one is relative stable and thus the protein stays in these conformations longer leading to a better CC, whereas structures far from the native one are less stable and more dynamic so that the AFM images reflect some averaged structures, resulting in poorer CCs. In a similar line, the probability distributions in the RMSD and the CC (Figure 5D, H, and L) suggest that there is the larger correlation between RMSD and CC as the best CC decreases; it also supports that structures far from the native one is more dynamic and thus can be fit with poor CCs.

## DISCUSSIONS

In this work, we modeled the AFM stage-protein interaction as a simple Lennard-Jones interaction that was uniformly applied on the xy-plane. In reality, the interaction between stage and specimen is more complicated. The surface of a real stage is actually composed of atoms. Thus, the stage potential would not be homogeneous in space. There is the electrostatic interaction between specimen molecules and stage atoms, depending on the chemical characteristics of the stage atoms. However, the interaction between stage and specimen has not been analyzed in detail and the shape of the stage strongly depends on the experimental setup and cannot be practically controlled in the atomic scale. Since the precise information is not available, it is not appropriate, at this stage, to introduce an extra complexity to a model. Therefore, here we employed the simplest interaction between the stage and the specimen. Because our biasing potential is independent from the stage potential, we can improve the stage potential with more accurate model of AFM stages, if necessary. Additionally, our method will be useful to investigate the precise model of AFM stage because one can optimize the parameter of AFM stage model and molecular structure simultaneously to make the difference between structure model and experimental AFM image as small as possible.

It is known that HS-AFM image contains an artifact called parachuting effect in some condition^5^. Although the recent research provides some solution on to this problem^33^, still it is good to deal with such a kind of errors. Two ways to work around this problem can be considered. The first one is to employ techniques accumulated in the field of image processing to remove such effects. The second is to change the formulation of our biasing potential to emulate parachuting effect. We used a symmetric gaussian function as a weighting function for XY-plane in this work for simplicity. However, it is possible to consider asymmetric spatial effects such as parachuting by changing weight asymmetrically according to the direction of scanning lines.

We also show that the effectiveness of our biasing potential strongly depends on the value of parameter κ. From our simulations, we conclude that the value from 0.5 to 1.5 is an appropriate range. It might depend on the shape of a target molecule and the size of the image to fit; therefore, some pre-sampling might be needed. To fit images that is considered to represent structures that are quite different from crystal structure, we may need to make our biasing potential stronger than the current cases. To automate the parameter tuning, the methods that are established for cryo-EM flexible fittings, e.g. changing potential strength dynamically, might be useful ^34^.

We tested the applicability of our method to experimental AFM images. While experimental images have an unpredictable noise, our method still found conformations that have high correlations with the experimental images. Especially for images that seem to change its conformation, however, some of the resulting conformations still look different from the corresponding experimental image. There can be several reasons. First, the shape of the cantilever tip may be less sharp than our expectation. To address this issue, we need to calibrate the value of parameter *σ* using experimental images that shows well predictable object. Next, there can be an artifact from the unavoidable noises in experimental images. To gain higher signal noise ratio, we can average several images as the current cryo-EM image analysis normally does. Another way is to use temporal information that AFM movie contains. Applying data-assimilation technique, we can use full information throughout the whole movie and achieve more accuracy. There is another information source, the experimental setup. AFM stage can have a charge distribution that modulates strongly to the orientation of the molecule. By estimating and applying the electrostatic potential between specimen molecules and the stage, the possible orientations and conformation will be restricted and then the resulting conformation will be more accurate.

Additionally, although we applied our novel biasing potential to coarse-grained molecular dynamics, it is potentially applicable to the all-atom fine-grained molecular dynamics simulation. Since the underlying method that generates pseudo-AFM images by combining smooth function works with atomic coordinates, there is no essential limitation to apply our biasing potential to the atomic models. Practically, because of the resolution of the available HS-AFM images, the atomic scale simulation might not be able to increase the quality of the structural model compared to the required computational efforts. But it is still interesting to run a short fine-grained molecular dynamics simulation with our biasing potential with the atomic structures that is reconstructed by structural modeling based on the result of our method.

## CONCLUSION

In this work, we proposed and tested the flexible fitting MD simulations with a biasing potential that quantifies the fitness to the experimental AFM image. We generated a smoothed form of the pseudo-AFM image generation method from a structure and a biasing potential that represents the correlation between the experimental and the structure-based AFM images. The biasing potential is differentiable so that we can apply it directly to MD simulations. By combining the AFM observations of biomolecules with MD simulation, our method would be a powerful tool that connects single molecule experiments with structural changes of proteins at work.

## Acknowledgement

The study was supported mainly by the Japan Science and Technology Agency (JST) grant JPMJCR1762 (ST), partly by the MEXT as “Priority Issue on Post-K computer” (ST), and by the RIKEN Pioneering Project “Dynamical Structural Biology” (ST). We appreciate Professor Noriyuki Kodera giving us AFM image data and for lessons on the machinery of AFM. We appreciate Suguru Kato for his technical supports and discussions. We thank Dr. Giovanni Brandani, Dr. Cheng Tan, Diego Ugarte and Hana Slevin for fruitful discussions. We also thank Shintaroh Kubo for giving a structure model of dynein and for discussions.

