## Supplemental Information for "Flexible Fitting of Biomolecular Structures to Atomic Force Microscopy Images via Biased Molecular Simulations"

**Rigid Body Metropolis Monte-Carlo Simulation**

While the flexible-fitting is the main theme of this work, for comparison we also performed the simulated annealing calculations for the rigid-body fitting to the AFM image. Here, we describe method details for the rigid-body fitting.

For this, Metropolis Monte-Carlo simulations with simulated annealing method was used to find the optimal conformation. Assuming that a target protein is placed on the stage plane, we consider only translations in xy-directions and rotations around x-, y-, and z- axes as trial movements. For the former, we drew random numbers from a uniform distribution in the range of $-0.01 nm <\Delta x (or \Delta y)<0.01nm$, and moved accordingly. For the rotation, in each cycle, we drew three angles from the uniform distribution in the range $-0.1 \pi<\Delta\theta<0.1\pi$, and rotated accordingly. In each of these translation or rotation moves, after the move, we placed the protein to be in contact with the stage at the bottom of the molecule. For each of these five trials, we either accept or reject the move by using the standard Metropolis criterion with $-5\sin^{-1} c.c.(\boldsymbol{R})$ as the "energy".

The correlation coefficient is the same as that in the flexible fitting. Explicitly,

$$c.c.(\boldsymbol{R}) = \frac{\sqrt{\sum_{p}^{pixels} H_{p}^{(exp)}H_{p}^{(sim)}(\boldsymbol{R})}}{\sqrt{\sum_{p}^{pixels} \left( H_{p}^{(exp)} \right)^{2}}\sqrt{\sum_{p}^{pixels} \left( H_{p}^{(sim)}(\boldsymbol{R}) \right)^{2}}}$$

Here, all the sums run over the pixels. The $H_{p}^{(exp)}$ represent the experimental AFM images, while $H_{p}^{(sim)}(\boldsymbol{R})$ is a pseudo-AFM image defined by eq.(1) for a current structure coordinate collectively represented as $\boldsymbol{R}$. It is demonstrated that essentially the same formula successfully works for cryo-EM-based flexible fitting problem^24,25^. In this study, we employ a "modified" form where the mean height of *H*_p_ is "not" subtracted both in the denominator and numerator of $c.c.(\boldsymbol{R})$. This is because the standard correlation coefficient does not change when $H_{p}^{(sim)}(\boldsymbol{R})$ is constantly shifted in z-direction. This is not a desired feature; we need to fit the absolute height (the z-coordinate), given that the stage corresponds to *z* = 0.

In the simulated annealing runs, the temperature is decreased exponentially in step with the initial and final temperatures, T = 1 and T = 0.0001, respectively. Each simulated annealing run contains 10^5^ MC steps.

As the reference image, we generated synthetic AFM image from randomly chosen configuration and applied spatially uncorrelated Gaussian noise on it with various value of the variance.

We performed 200 independent runs for dynein and 100 for hemoglobin.
